# Do pollinators play a role in shaping the essential amino acids found in nectar?

**DOI:** 10.1101/2024.09.30.615756

**Authors:** Rachel H. Parkinson, Eileen F. Power, Kieran Walter, Alex E. McDermott-Roberts, Jonathan G. Pattrick, Geraldine A. Wright

## Abstract

- Plants produce floral nectar as a reward for pollinators, which contains carbohydrates and amino acids (AAs). We designed experiments to test whether pollinators could exert selection pressure on the profiles of AAs in nectar.
- We used HPLC to measure the free amino acids and sugars in the nectar of 102 UK plant species. Six distinct profiles of essential amino acids (EAAs) were defined using the relative proportions of AAs with a clustering algorithm; we then tested bumblebee (*Bombus terrestris*) preferences for the EAA profiles and proline using a two-choice assay.
- We found a phylogenetic signal for the proportions of phenylalanine, methionine and proline as well as the total concentrations of essential and non-essential AAs. However, there was no phylogenetic signal for EAA profile. Bumblebees did not exhibit a preference for any of the six EAA nectar profiles, however, four of the EAA profiles stimulated feeding. In contrast, bumblebees avoided proline in an inverse concentration-dependent manner.
- Our data indicate that bees are likely to have mechanisms for the post-ingestive evaluation of free AAs in solution but are unlikely to taste EAAs at nectar-relevant quantities. We predict that EAAs increase nectar value to bumblebees post-ingestively.

## Introduction

Floral nectar is produced by plants to attract animals to flowers for pollination. Most floral nectar is dominated by the presence of sugars (sucrose, glucose, and fructose), but the second most abundant metabolites are free amino acids (AAs) (Nicolson & Thornburg, 2007). Free AAs are ubiquitous in nectar, and although quantities can vary considerably (Baker & Baker, 1975; Gottsberger *et al*., 1984; Gardener & Gillman, 2001a; Vandelook *et al*., 2019), they are present at concentrations (micromolar to millimolar) that are substantially lower than the carbohydrates (Nicolson, 2022).

Free AAs in nectar can come from the phloem but can also be produced by the nectary itself (reviewed in Göttlinger & Lohaus, 2024). The concentrations of free AAs in nectar are typically lower than in other plant tissues and may be as much as 100-fold lower than in the nectaries and phloem (Lohaus & Schwerdtfeger, 2014; Bertazzini & Forlani, 2016; Göttlinger & Lohaus, 2024). While the nectar AA profile (relative abundance of the different AAs) may resemble that in the nectaries (Göttlinger & Lohaus, 2022) it often differs from nectary (Göttlinger & Lohaus, 2024) and phloem (Lohaus & Schwerdtfeger, 2014; Bertazzini & Forlani, 2016) composition, suggesting that the nectar AA profile is more than just a simple filtering of phloem constituents. A hypothesis driving this behaviour is that nectar amino acids are driven by pollinator selection (Tiedge & Lohaus, 2017; Göttlinger *et al*., 2019).

Several authors have shown associations between pollinator groups and nectar AA profile and concentration (Baker & Baker, 1975; Petanidou *et al*., 2006; Tiedge & Lohaus, 2017; Göttlinger & Lohaus, 2024) though in some situations, AAs do not appear to be as important as other nectar components (Göttlinger *et al*., 2019; Vandelook *et al*., 2019). One criticism of looking for correlations between nectar AAs and pollinator group is that AA concentrations can vary considerably among individuals within a plant species (Gardener & Gillman, 2001a; Gijbels *et al*., 2015b). However, large scale studies have found that while concentration varies, the relative abundance of individual AAs is much more consistent within species (Gardener & Gillman, 2001a) suggesting AA profile may be a better target for pollinator-driven selection than absolute concentration. Pollinator discrimination between nectars based on AA composition could reflect metabolic demand (Mevi-Schütz & Erhardt, 2005) or taste (Gardener & Gillman, 2002).

Insect pollinators, such as bees, rely on pollen as their primary protein source (Wright *et al*., 2018) and only secondarily derive benefits from nectar sources of free AAs (Nicolson, 2022). Other pollinators that do not consume pollen like butterflies may depend more on nectar-derived amino acids (Erhardt & Rusterholz, 1998; Mevi-Schütz & Erhardt, 2005; Beck, 2007). For this reason, one might predict that bees do not exhibit strong preferences for nectar solutions containing free AAs. However, honeybees have preferences for sugar solutions that contain one of the EAAs over solutions containing one of the non-essential AAs (NEAAs), with the strongest preference observed towards phenylalanine (Hendriksma & Shafir, 2016), a compound frequently found in floral nectar in high concentrations (Petanidou *et al*., 2006). Other free AAs in sugar solutions, such as methionine, have been reported to suppress feeding in honeybees (Inouye & Waller, 1984; Simcock *et al*., 2014). While many studies have examined individual AAs in nectar, none have reported how the profiles or mixtures of AAs found naturally occurring in nectar influence the preferences of pollinators.

Some NEAAs are also important to pollinators. For example, proline is the dominant amino acid in the haemolymph of both honeybees (Crailsheim & Leonhard, 1997) and bumblebees (Stabler *et al*., 2015). Proline can also act as a substrate for powering insect flight (Bursell, 1975; Auerswald & Gäde, 1999), though evidence for this role in bees is more equivocal. In honeybees, haemolymph proline concentration decreases significantly after flight (Barker & Lehner, 1972; Micheu *et al*., 2000), though the proportional contribution compared to carbohydrates is minimal (Barker & Lehner, 1972). Isolated flight muscles from *Bombus impatiens* bumblebees showed significant increases in the rate of respiration when exposed to proline (Teulier *et al*., 2016); however, this effect was not found with mitochondria from males of the bumblebee *B. terrestris* (Syromyatnikov *et al*., 2013). More recent work with whole living bumblebees suggests that, at least in *Bombus impatiens*, proline is most likely used as a sparker for carbohydrate metabolism for the first few minutes of flight (Stec *et al*., 2021). Honeybees appear to prefer sucrose solutions containing 2–10 mM proline concentrations (Carter *et al*., 2006; Bertazzini *et al*., 2010) but find higher proline concentrations aversive (Carter *et al*., 2006; Simcock *et al*., 2014). Other bee species may have different preferences, however. One study reported that nectar-relevant proline of ∼2 mM appeared to suppress sugar solution consumption in bumblebees (Bogo *et al*., 2024).

In this study, we analysed the carbohydrate and amino acid profiles of nectar from 102 plant species from the UK and tested whether these metabolites reflected phylogenetic relationships among the selected plant species. Based on previous work where we observed that the EAAs were the most important components of protein for bumblebees (Stabler *et al*., 2015), we examined the impact of EAAs on bumblebee preferences for nectar. Using a k-means clustering algorithm, we identified six distinct essential amino acid (EAA) profiles in UK plant species nectar. Buff-tailed bumblebees were given a choice of the solutions over 24 h but did not exhibit preferences or aversions towards the solutions containing the EAAs. While the bees did not distinguish between the different solutions, they consumed more overall when certain EAA profiles were present. Additionally, we examined the effects of proline in nectar and found that proline influenced feeding preferences in a concentration-dependent manner. These findings demonstrate that the profile of EAAs at nectar-relevant concentrations are valuable to bumblebees but do not affect their initial preferences, indicating that they do not rely on their sense of taste to detect them in solution. Our results suggest that floral nectar is under selective pressure from pollinators. EAAs increase nectar value in a way that is independent of sensory cues.

## Materials and Methods

### Nectar data collection

Nectar data was collected in 2011 and 2012 from sites in north-eastern and southern England. We sampled nectar using 1 µl microcapillary tubes (Hirschmann Laborgeräte GmbH & Co. KG, Eberstadt, Germany) to extract raw nectar from open flowers with healthy appearances (no signs of senescence). The microcapillary method minimises potential contamination from e.g. pollen or vascular fluid (Power *et al*., 2018). The volume of nectar obtained from individual flowers was measured in the field or laboratory based on the length filled of the microcapillary tube. Nectar samples were then diluted with UHPLC gradient grade water (Fisher Scientific UK Ltd., Loughborough, United Kingdom) to meet minimal sample volume requirements for carbohydrate and amino acid analyses. The optimal dilution of nectar:water needed was 1:2000, requiring approximately 0.05 µl of raw nectar (to make 100 µl of solution). Samples were centrifuged to remove any residual plant material.

### Nectar sample analysis

Nectar sugar composition (sucrose, glucose, and fructose) was analysed using high performance ion chromatography (HPIC). For each sample, approximately 30 µl of diluted nectar was inserted into an analysis vial to ensure optimal immersion of the autosampler syringe, and 20 µl of this nectar sample was injected via a Rheodyne valve onto a Carbopac PA-100 column (Dionex, Sunnyvale, California, USA) fitted with a Dionex Carbopac PA-100 BioLC guard (4 x 50 mm). Sample components were eluted isocratically using 100 mM NaOH (de-gassed by helium gas) flowing at 1 ml min^-1^ for 10 min at room temperature (RT).

We recorded the chromatographic profiles using pulsed amperometric detection with an ED40 electrochemical detector (Dionex, Sunnyvale, California, USA), and analysed the elution profiles using Chromeleon v 6.8 (Thermo Fisher Scientific Inc., MA, USA). The HPIC was calibrated at least twice every 24 h period for all compounds of interest by injecting calibration standards with concentrations of 10 ppm each. Standard solutions were made from the solid forms of each sugar (Sigma-Aldrich, St. Louis, MO, USA). Measured concentrations were rescaled to the original concentration using the dilution factor.

Amino acid composition was determined using ultra high-performance liquid chromatography (uHPLC). This gave the concentrations of all 20 standard protein-forming amino acids and GABA. We used an automated pre-column derivatization programme for the autosampler (Ultimate 3000 Autosampler, Dionex, Thermo Fisher Scientific Inc.) immediately before injection to pre-treat 10 µl of sample. This was comprised of: 1 min with 15 µl of 7.5 mM o-phthaldialdehyde (OPA) and 225 mM 3-mercaptopropionic acid (MPA) in 0.1 M sodium tetraborate decahydrate (Na_2_B_4_O_7_.10H_2_O), pH 10.2; 1 min with 10 μl of 96.6 mM 9-fluroenylmethoxycarbonyl chloride (FMOC) in 1 M acetonitrile; followed by the addition of 6 μl of 1M acetic acid. After pre-treatment, 30 µl of the amino acid derivatives were injected onto a 150 x 2.1 mm Accucore RP-MS uHPLC-column (Thermo Fisher Scientific Inc.). Elution solvents were: A = 10 mM di-sodium hydrogen orthophosphate (Na_2_HPO_4_), 10 mM Na_2_B_4_O_7_.10H_2_O, 0.5 mM sodium azide (NaN_3_), adjusted to pH 7.8 with concentrated HCl, and B = Acetonitrile/Methanol/Water (45/45/10 v/v/v). Elution of the column occurred at a constant flow rate of 500 μl min^-1^ using a linear gradient of 3 to 57% (v/v) of solvent B over 14 min, followed by 100% solvent B for 2 min and a reduction to 97% solvent B for the remaining 4 min (Power *et al*., 2018).

The derivatives were detected by fluorescence (Ultimate 3000 RS Fluorescence Detector, Dionex, Thermo Fisher Scientific, OPA: excitation at 330 nm and emission at 450 nm, FMOC: excitation at 266 nm and emission at 305 nm) and quantified by automatic integration after calibration of the system with amino acid standards. Reference calibrations (for all amino acids) were conducted following the processing of each batch of 20 samples by injecting calibration standards. Elution profiles were analysed using Chromeleon software v. 6.8 (Thermo Fisher Scientific Inc). Amino acid peaks were detected automatically based on pre-calibrated elution times, with all peaks checked to ensure correct identification.

### Nectar dataset analyses

Concentrations of nectar sugars were adjusted based on sucrose-equivalency, such that for glucose and fructose 1 M = 0.5 sucrose-equivalent molarity (se-M, Fleming et al., 2008; Leseigneur & Nicolson, 2009). We summed the se-M concentrations to obtain a measure of total sugars and calculated the proportion of the total represented by each sugar. For amino acids, the proportions of the essential amino acids (EAAs: His, histidine; Thr, threonine; Arg, arginine; Val, valine; Met, methionine; Trp, tryptophan; Phe, phenylalanine; Ile, isoleucine; Leu, leucine; and Lys, lysine) and non-essential amino acids (NEAAs: Asn, asparagine; Asp, aspartic acid; Ser, serine; Gln, glutamine; Gly, glycine; Ala, alanine; GABA gamma-aminobutyric acid; Tyr, tyrosine; Cys, cystine; and Pro, proline) were calculated as proportions of the total amino acids (the summed concentrations of all essential and non-essential amino acids). Data were averaged across samples for each species.

To visualise the high-dimensional relationships between nectar EAA profiles across species, we employed t-distributed Stochastic Neighbour Embedding (t-SNE). We standardised (z-scored) the EAA proportion data, and the perplexity parameter was set to 20 to balance local and global aspects of the data. We determined the optimal number of clusters in the t-SNE reduced-dimensional space with a Silhouette Analysis, which measures the cohesion and separation of clusters. The average silhouette width was calculated for k = 2 to 20, and the number of clusters with the highest silhouette width was selected as optimal. Using this optimal k, we applied K-means clustering to the t-SNE coordinates. We reconstructed the proportions of each EAA (relative to total amino acids) and sucrose-equivalent sugar proportions for sucrose, glucose, and fructose from the nectar clusters. The mean nectar concentrations of EAAs and sugars were also reconstructed from the species in each cluster.

We assembled a phylogenetic tree by pruning an existing dated tree with the species from our dataset (Zu *et al*., 2021). We dealt with missing species in the following ways: 1) if the missing species was the only representative of a genus, we replaced it with another species from the same genus; and 2) species were removed if they were not in the template tree and other representative species from the same genus were present.

### Preference assays

We assessed the preferences of bumblebees (*Bombus terrestris audax*) for sugar mixtures with or without amino acids using a preference assay. Bumblebees were reared by Biobest (Westerlo, Belgium), and purchased from Agralan (Ashton Keynes, UK), and maintained at RT in their colony boxes with access to Biogluc (Biobest, Westerlo, Belgium) and provided with pollen. For the preference assays, worker (female) bees were caught while exiting the colonies and assigned to treatment groups such that at least two colonies were tested in every group. For the duration of the assays, bees were housed in groups of five and provided with *ad libitum* access to the test solutions from four modified Eppendorf tubes (two on either side of the cages).

We performed three separate preference assays to test the bees’ preferences for EAAs and proline: 1) comparing the sugar backgrounds with and without the mean nectar EAA concentrations for each of the six clusters identified from the clustering analysis (Table 1); 2) a concentration gradient of cluster B EAA and sugar profiles at 1 to 1000 μM versus the sugar background alone; and 3) a concentration gradient of proline (1 to 1000 μM) versus the sugar background containing 0.5 M each of glucose and fructose. Test solutions were prepared with reagent grade sucrose, glucose, fructose, essential amino acids, and proline (Merck Life Science, UK).

**Table 1:**
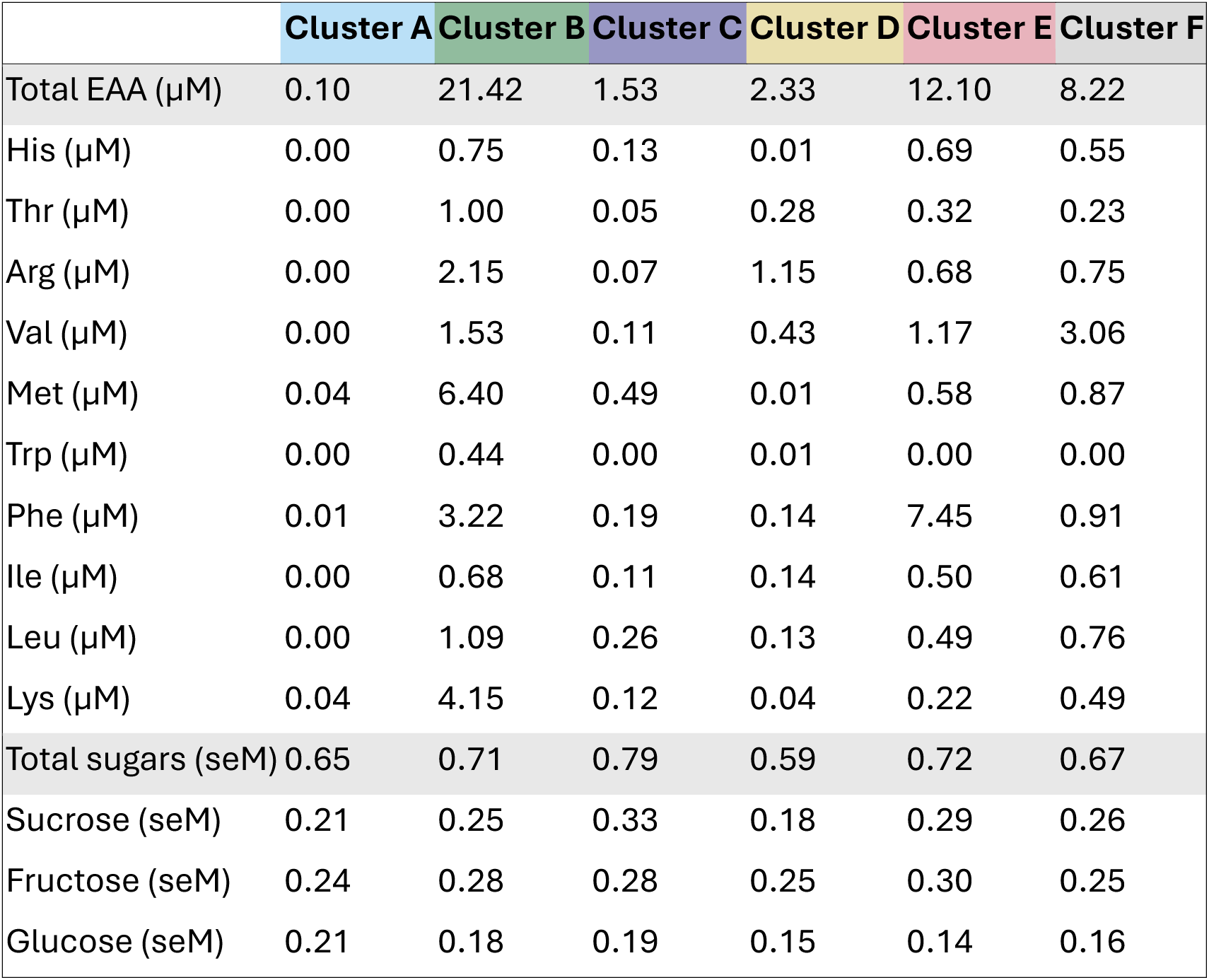
Amino acid profiles of clusters from nectars dataset. Values are given as mean concentrations for each cluster, with sugar concentrations reported as sucrose-equivalent Molarity (seM).

|  | Cluster A | Cluster B | Cluster C | Cluster D | Cluster E | Cluster F |
| --- | --- | --- | --- | --- | --- | --- |
| Total EAA (μM) | 0.10 | 21.42 | 1.53 | 2.33 | 12.10 | 8.22 |
| His (μM) | 0.00 | 0.75 | 0.13 | 0.01 | 0.69 | 0.55 |
| Thr (μM) | 0.00 | 1.00 | 0.05 | 0.28 | 0.32 | 0.23 |
| Arg (μM) | 0.00 | 2.15 | 0.07 | 1.15 | 0.68 | 0.75 |
| Val (μM) | 0.00 | 1.53 | 0.11 | 0.43 | 1.17 | 3.06 |
| Met (μM) | 0.04 | 6.40 | 0.49 | 0.01 | 0.58 | 0.87 |
| Trp (μM) | 0.00 | 0.44 | 0.00 | 0.01 | 0.00 | 0.00 |
| Phe (μM) | 0.01 | 3.22 | 0.19 | 0.14 | 7.45 | 0.91 |
| Ile (μM) | 0.00 | 0.68 | 0.11 | 0.14 | 0.50 | 0.61 |
| Leu (μM) | 0.00 | 1.09 | 0.26 | 0.13 | 0.49 | 0.76 |
| Lys (μM) | 0.04 | 4.15 | 0.12 | 0.04 | 0.22 | 0.49 |
| Total sugars (seM) | 0.65 | 0.71 | 0.79 | 0.59 | 0.72 | 0.67 |
| Sucrose (seM) | 0.21 | 0.25 | 0.33 | 0.18 | 0.29 | 0.26 |
| Fructose (seM) | 0.24 | 0.28 | 0.28 | 0.25 | 0.30 | 0.25 |
| Glucose (seM) | 0.21 | 0.18 | 0.19 | 0.15 | 0.14 | 0.16 |

The assays ran for 72 hours (n = 15 cages per group). During the first 48 hours, bees were fed with the sugar background solutions only (no amino acid choice solutions). We used the data from the second 24 hours (after a 24-hour acclimation period) as a no-choice control to demonstrate whether the bees preferred to drink from either side of the cage and obtain baseline consumption volumes. For the final 24-hours, bees were given a choice between the sugar solution, or a solution containing EAAs in the same sugar background.

We recorded the volume consumed by weighing the solutions before and after each 24-hour period, with the average evaporation quantities taken from the totals. A preference index was calculated using the following formula:

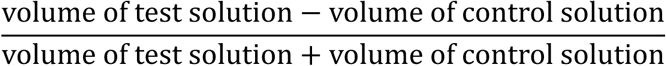

We recorded any mortality on each day of the experiment and scaled the average consumption per bee by the number of surviving individuals. Following each assay, bees were euthanised by freezing at -20°C.

To estimate the proportion of the 102 plant species which are visited by bumblebees, we used The Database of Pollinator Interactions (Balfour *et al*., 2022). A species was classed as visited by *Bombus terrestris/lucorum* if there were any recorded interactions from bees in these two groups respectively. *Bombus terrestris* and *Bombus lucorum* workers are indistinguishable in the field and so typically grouped together.

#### Statistical analyses

All statistical analyses were performed in R version 4.3.3 (R Core Team, 2024). Data were tested for normality with Shapiro-Wilk tests. Data that were not normally distributed were transformed or tested with non-parametric approaches. We assessed whether the average concentrations of amino acids (log_10_ concentrations of EAAs and proline) or sugars varied across species using non-parametric Friedman tests, accounting for repeated measures within species. Pairwise comparisons were conducted using the Nemenyi-Wilcoxon-Wilcox all-pairs *post hoc* test, with a single-step p-value adjustment for multiple comparisons. Amino acid proportion data was logit transformed, following which we explored the correlation between the average proportions of EAAs and proline across species using Pearsons’s correlation coefficients, retaining only significant correlations (p < 0.05) for interpretation and visualisation.

We compared amino acid and sugar profiles across cluster assignments using PERMANOVA with adonis post hoc multiple comparisons performed with the ‘vegan’ package (Oksanen J *et al*., 2024). We fitted beta regression generalised mixed effects models (GLMM) with a logit link function to assess differences in the proportions of amino acids and sugars within each cluster, accounting for repeated measures by including the species as a random effect. The models were fitted using the ‘glmmTMB’ package in R (Brooks *et al*., 2017). We assessed which amino acids represented the largest proportions in each cluster by comparing the marginal means with the overall mean using the ‘emmeans’ package (Russell *et al*., 2021). We assessed the phylogenetic signal of the nectar profiles (the 2 dimensions from the t-SNE analysis, individual amino acid proportions, total amino acid concentrations, sugar concentrations, and sugar proportions) with Blomberg’s K and Pagel’s Lambda (M. Pagel, 1999; Blomberg *et al*., 2003) using the ‘phylosignal’ package in R (Keck *et al*., 2016).

For the behavioural data, we modelled the preference indices and total volume consumed using linear mixed effects models (LME) comparing between the sugar only (no choice) and test days, as well as across groups, with cage ID as the random effects using the ‘lmerTest’ package in R (Kuznetsova *et al*., 2017). Post hoc tests were performed with the emmeans, with p-values adjusted for multiple comparisons using the Tukey method. Mortality was assessed over the duration of each preference assay using Kaplan-Meier survival analysis. Exact p-values were provided unless p < 0.0001.

## Results

### Nectar amino acid and sugar concentrations

Our nectar dataset included 102 plant species, 53% of which are known to be pollinated by *Bombus terrestris,* and 57% by other bumblebee species (Balfour et al., 2022, SI Table 1). However, 86% of the genera included have at least one species pollinated by bumblebees (Balfour et al., 2022, SI Table 1). We used uHPLC and HPIC to quantify the concentrations of amino acids and sugars, respectively, in the nectar of 102 species across 78 genera and 36 families (SI Table 1). The nectar showed substantial variation in total EAA concentration (mean = 9.16 µM, sd = 22 µM) and NEAA concentration (mean = 5.94 µM, sd = 13.6 µM), with a range spanning nanomolar to millimolar concentrations across species (Figure 1A). The concentrations of individual AAs also varied considerably (Figure 1B). Trp exhibited the lowest mean concentration in nectar (mean = 0.113 µM; sd = 0.581 µM) whereas Pro (mean = 217 µM, sd = 124 µM) was almost always present at higher concentrations than any of the AAs. The most extreme value for proline was observed in the nectar of *Iris pseudocorus* (mean = 11.4 mM).

**Figure 1:**
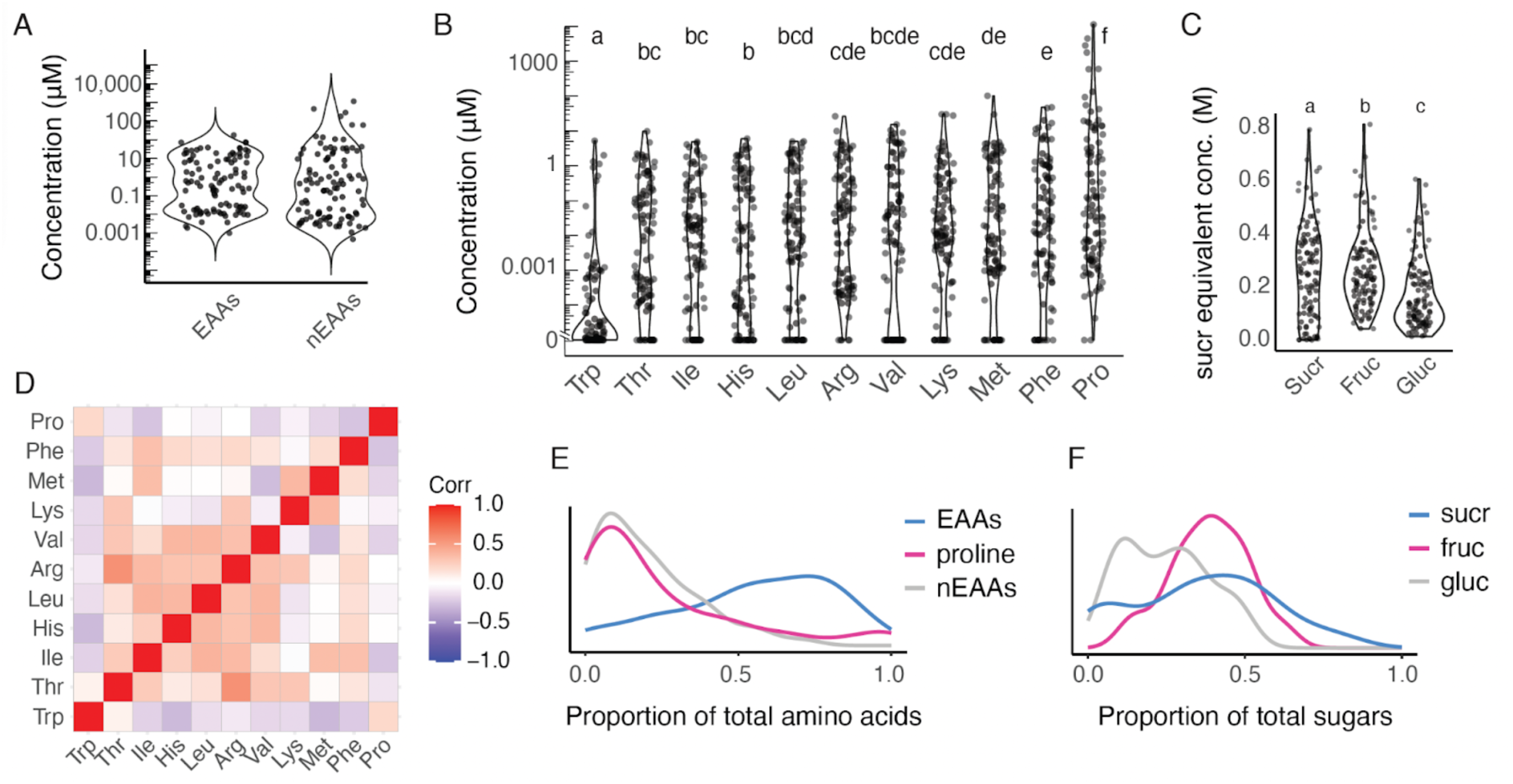
Amino acids and nectar sugars from 102 plant species. A: Concentrations of essential amino acids (EAAs) and non-essential amino acids (nEAAs) for all species (n = 102). Datapoints represent average concentrations per species (n = 102 species, min = 3, max = 28 samples/species). B: Nectar concentrations of EAAs and proline (Friedman χ^2^ = 277, df = 10, p < 0.0001, n = 102). Significant differences between groups (Nemenyi-Wilcoxon-Wilcox post hoc) are denoted with letters. Datapoints represent average concentrations per species. C: Sucrose equivalent nectar concentrations of sucrose, fructose, and glucose (Friedman χ^2^ = 51.8, df = 2, p < 0.0001, n = 102). Significant differences between groups (Nemenyi-Wilcoxon-Wilcox post hoc) are denoted with letters. Datapoints represent average concentrations per species. D: Pearson correlation coefficients of the logit-transformed proportions of EAAs and proline across all species in the dataset. Only significant correlations are plotted (non-significant correlations are set to 0). E: The distribution of the average essential amino acids (EAAs), proline, and other non-essential amino acids (nEAAs) as a proportion of total amino acids across all species. F: Distribution of the average sucrose equivalent concentrations of sucrose (sucr), fructose (fruc), and glucose (gluc) across all species.

We used an amino acid concentration cutoff of 0.001 μM to quantify which amino acids were represented in the species in our nectar dataset. Pro was found in the highest proportion of any AA (87% of species). Of the other NEAAs, Asn, GABA, and Gln were present in only approximately 20% of species, while Asp, Glu, Ser, Gly, Ala, Tyr, and Cys were present in approximately 60% of species. Each of the EAAs were present in between 54% (His and Thr) and 76% (Phe) of species, except for Trp, which was found in just 17% of species at concentrations > 0.001 μM. When looking at the number of EAAs found in individual species, 32% of species had fewer than five EAAs, while 47% had between eight and 10 EAAs. Patterns were similar for NEAAs, with 40% of species having fewer than five NEAAs, and 42% with between eight and 11 NEAAs.

The three nectar sugars also varied, with significantly lower sucrose-equivalent molarity (se-M) concentrations of glucose (mean = 0.17 M, sd = 0.13 M) compared to fructose (mean = 0.27 M, sd = 0.15 M) or sucrose (mean = 0.26 M, sd = 0.18 M, Figure 1C). The proportions of some individual EAAs + Pro were correlated across samples, with the highest positive correlation between Arg and Thr, and the largest negative correlation between Met and Trp (Figure 1D). On average, EAAs represented a greater proportion of the total amino acids, but some species had nectar containing almost exclusively proline (Figure 1E). Fructose was the dominant nectar sugar in most species, and no species had glucose-dominated nectar (Figure 1F).

### Clustering analysis of essential amino acid profiles

To identify distinct patterns of nectar composition based on the sugars and EAAs present in our samples, we used t-distributed stochastic neighbour embedding (t-SNE) with k-means clustering (Figure 2A-B). This clustering revealed six distinct nectar types (PERMANOVA: F_5_ = 27.6, R^2^ = 0.59, p = 0.001) driven by the proportions of individual EAAs (Figure 2C). Within each cluster, one or two EAAs were present at significantly higher proportions than the other EAAs: for example, Met and Lys (cluster A), Arg and Met (cluster B), Met and Leu (cluster C), Thr and Arg (cluster D), Val and Phe (cluster E), and Val (cluster F).

**Figure 2:**
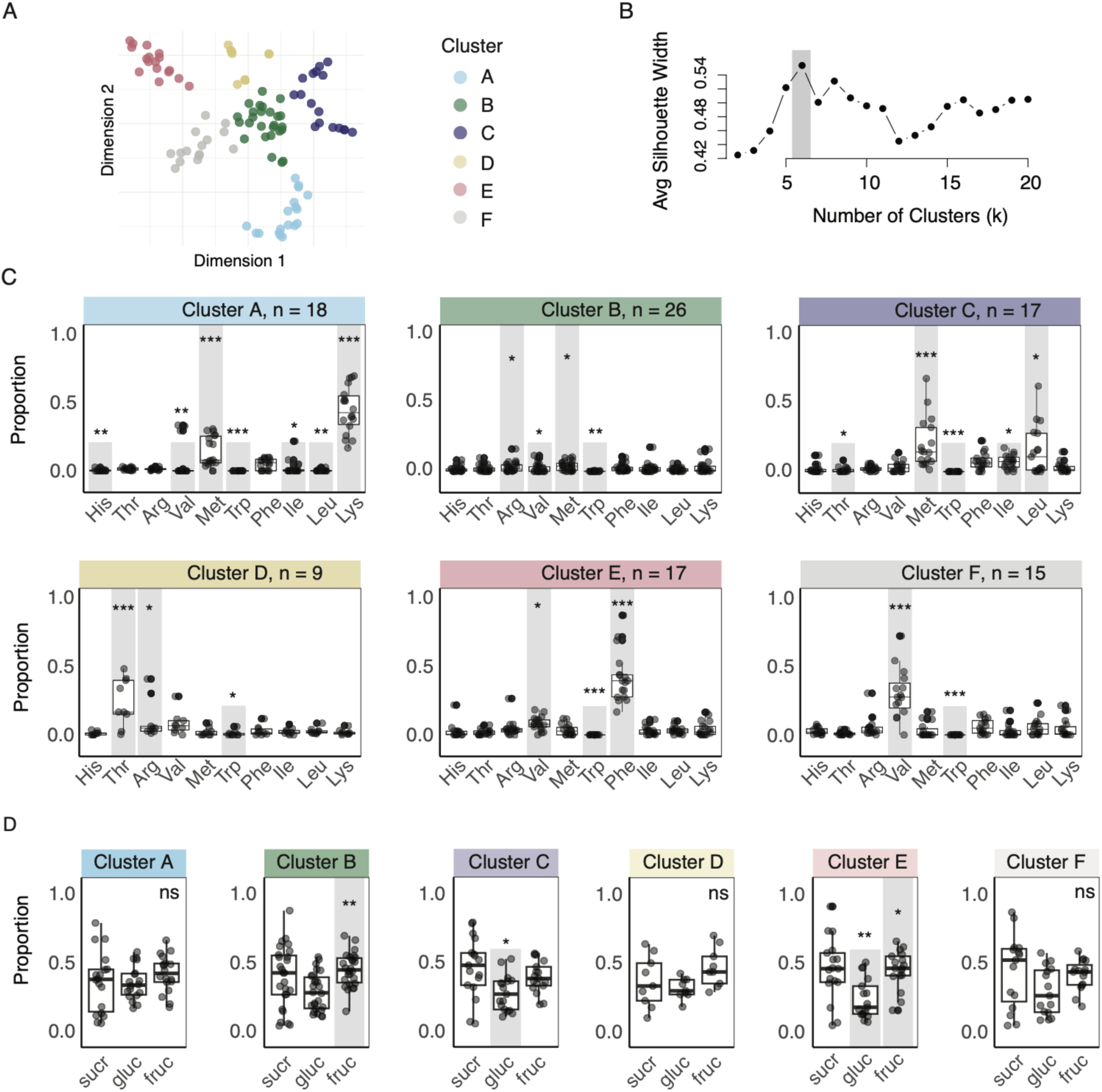
Clustering of the proportion of total amino acids represented by each EAA. A: t-SNE dimensionality reduction of the EAA proportion data. Each point represents a species, and the distance between points reflects the similarity of EAA profiles (n = 102). Clustering with k-means revealed 6 groupings (coloured) based on EAAs. B: Silhouette plot showing silhouette widths against the number of clusters. K = 6 was identified as the optimal cluster number with the highest silhouette width. C: EAA profiles varied significantly across clusters (PERMANOVA: F_5_ = 27.6, R^2^ = 0.59, p = 0.001), with significant differences in amino acid profiles between each cluster (adonis pairwise multiple comparisons, adjusted p = 0.015). Within each cluster, the relative proportions of EAAs and proline varied (GLMM, cluster A: χ^2^ = 330, df = 9, p < 0.0001; cluster B: χ^2^ = 52.6, df = 9, p < 0.0001; cluster C: χ^2^ = 89.6, df = 9, p < 0.001; cluster D: χ^2^ = 53.0, df = 9, p < 0.0001; cluster E: χ^2^ = 344, df = 9, p < 0.0001; cluster F: χ^2^ = 94.1, df = 1, p < 0.0001). Amino acids with significantly higher or lower estimated marginal means compared to the overall mean of the cluster are highlighted with long or short grey bars, respectively. Asterisks denote significance of post hoc tests. D: Sugar profiles did not vary significantly across clusters (PERMANOVA: F_5_ = 0.805, R^2^ = 0.0402, p = 0.59). Within cluster, there were significant differences between the proportions of sugars for clusters B, C, and E (GLMM, cluster B: χ^2^ = 9.79, df = 2, p = 0.0075; cluster C: χ^2^ = 6.19, df = 2, p = 0.045; cluster E: χ^2^ = 10.6, df = 2, p = 0.0049). There were no significant differences between the proportions of sugars in clusters A, D and F (cluster A: χ^2^ = 3.48, df = 2, p = 0.18; cluster D: χ^2^ = 5.77, df = 2, p = 0.056; F: χ^2^ = 2.90, df = 2, p = 0.23). Sugars with significantly higher or lower estimated marginal means compared to the overall mean of the cluster are highlighted with long or short grey bars, respectively. Asterisks denote significance of post hoc tests (emmeans).

The clusters also displayed distinct total concentrations of EAAs (Table 1), ranging from 0.1 µM (cluster A) to 21.4 µM (cluster B). The proportions of nectar sugars, however, did not vary significantly across clusters (PERMANOVA: F_5_ = 0.805, R2 = 0.0402, p = 0.59), although there were differences in the proportions of sugars within some of the clusters (Figure 2D, Table 1).

We compared whether the cluster assignments were predictive of whether the species of plants were known to be pollinated by bumblebees (Balfour *et al*., 2022). There were differences in the proportions of species associated with *Bombus terrestris*: only 33% of the species in cluster A are known to be visited by *Bombus terrestris,* while 78% of the species in cluster B have been associated with *Bombus terrestris* visitors (Balfour *et al*., 2022), however these differences were not significant (GLM, Χ2 = 5.37, df = 5, p = 0.373).

### Phylogenetic signal in nectar composition

The presence of phylogenetic signal was quantified using Blomberg’s K and Pagel’s Lambda (M. Pagel, 1999; Blomberg *et al*., 2003). These analyses revealed significant phylogenetic signal across several measures, most notably for the total concentration of EAAs, where there was a strong signal (λ = 0.913, p = 0.001, K = 0.44, p = 0.009, SI Figure 1, SI Table 2). We also found weaker signal for the total concentration of NEAAs (SI Figure 1A), for the proportions of Met, Phe, and Pro (Figure 3, SI Table 2) and the proportions of glucose and sucrose in nectar (SI Figure 1B); though each of these parameters were only significant from one of the two indices. Pagel’s Lambda was significant for the proportion of Met, indicating that closely related species have more similar Met profiles. In contrast, Blomberg’s K was significant for the proportions of Pro and Phe. None of the other EAAs showed any significant phylogenetic signal (SI Table 2). Despite the strong signal for the total concentration of EAAs, we found negligible signal for the profile of EAAs (as quantified by the t-SNE dimensionality reduction). This suggests that while total amino acid concentration and individual EAAs may be determined by phylogeny, the overall blend of AAs may be driven by other factors such as pollinator preference.

**Figure 3:**
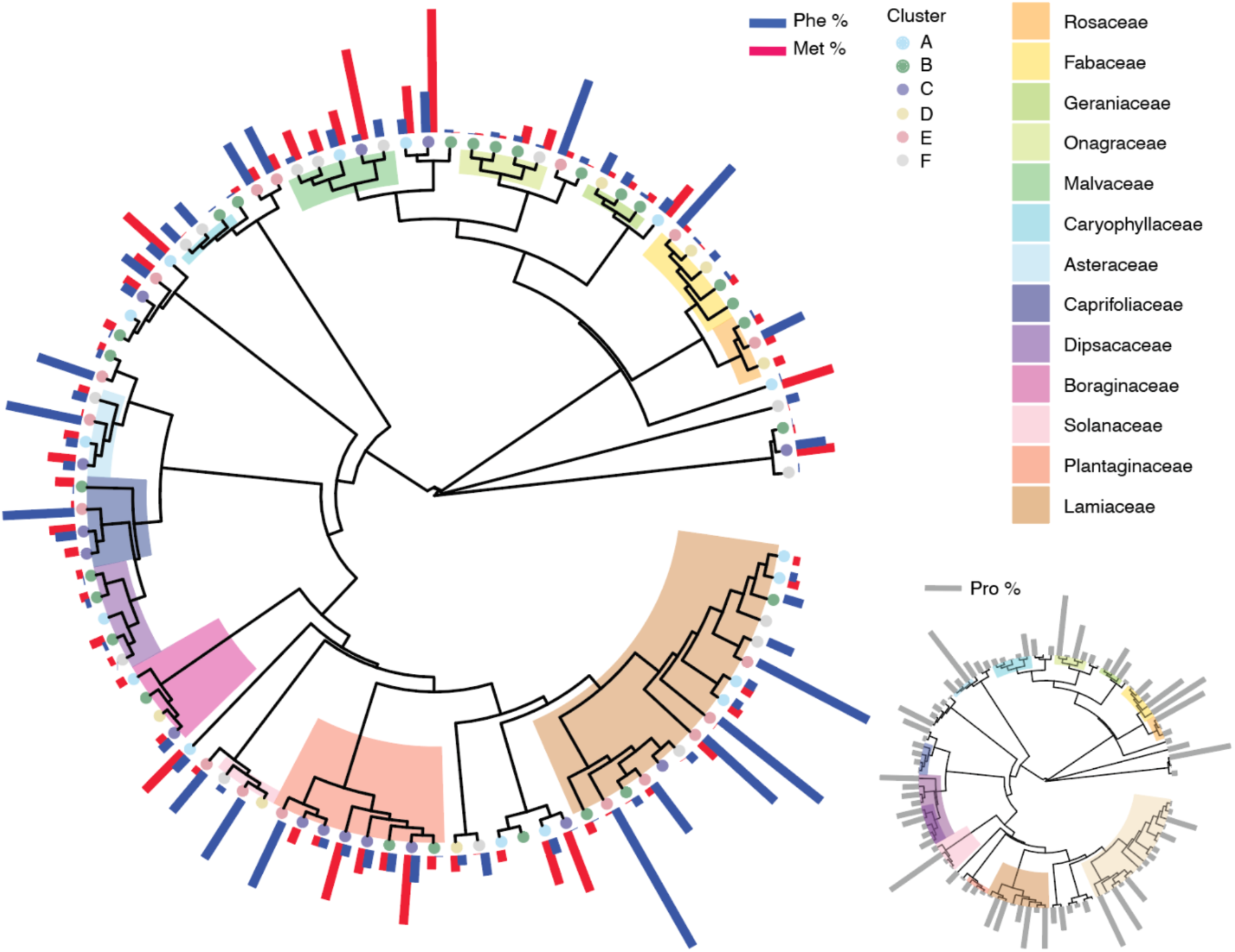
Phylogenetic analysis of amino acid profiles. A: Dated phylogenetic tree containing 96 species from our nectar dataset (pruned from Zu et al 2021). Families with at least three representative species are labelled with coloured blocks. Tip point circles show the cluster assignment based on the proportions of EAAs. Bars show the proportions of phenylalanine (Phe %) and methionine (Met %) for each species, as these EAAs had a phylogenetic signal. The inset phylogenetic tree shows the proportion of proline (Pro %) across species, which was the only non-EAA with a phylogenetic signal.

### Bumblebees drink more when EAAs are present in feeding choices

We created nectar solutions based on the nectar EAA profiles returned by the cluster analysis using the mean nectar concentrations of EAAs and sugars from each cluster (Table 1). In all cases, the bees did not show a preference for nectar sugar mixtures containing EAAs over the sugar mixtures alone (Figure 4A), indicating that EAA profiles did not drive an aversion to or a preference for nectar. However, if the total volume of solution consumed during the test period is considered, we found that for four out of the six nectar solutions, the bees consumed more food overall when EAAs were present in one of the solutions at nectar relevant concentrations (Figure 4B).

**Figure 4:**
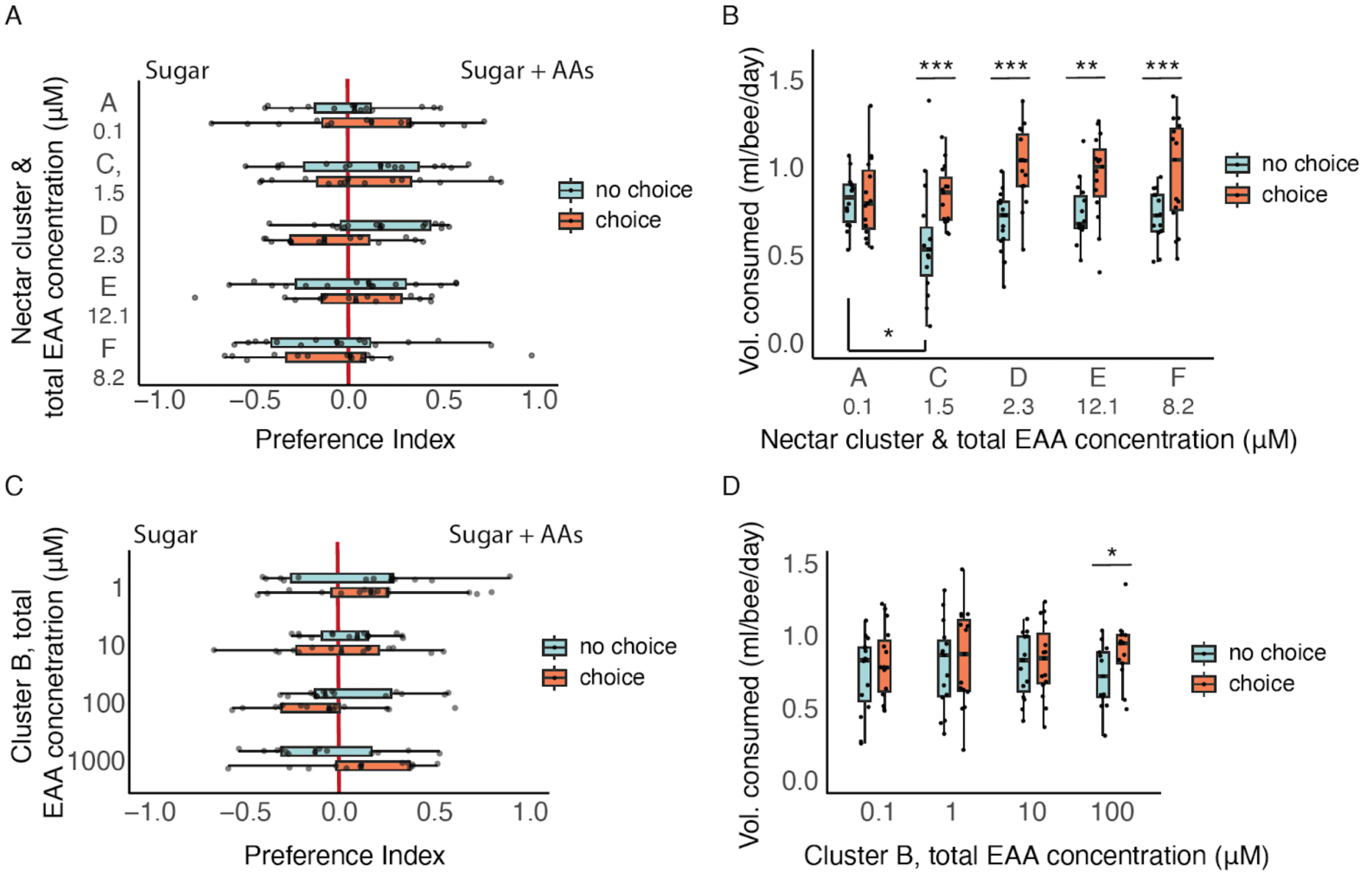
Bumblebee preferences for sugar solutions with nectar-relevant amino acid profiles. A: Preference assay using sugar and amino acid profiles from clustering analysis at nectar-relevant concentrations (see Table 1, n = 15 cages per group). Total EAA concentrations for each cluster are shown on the y-axis. There was no significant effect of adding EAAs on preferences (LME: choice, F_1. 143_ = 0.443, p = 0.51; cluster, F_4. 143_ = 0.826, p = 0.51). Boxplots show preferences on acclimation day (no choice) and test day (choice), with points representing individual cages. B: Total volume consumed (summing sugar-only and amino acid solutions from A, n = 15 cages per group). Significant interaction between day and nectar group (LME, day*group: F_4. 69_ = 3.36, p = 0.014, day: F_1, 69_ = 52.8, p < 0.0001; nectar group: F_4, 69_ = 1.76, p = 0.15). Asterisks indicate significant post hoc comparisons (emmeans, Tukey adjustment). C: Preference assay comparing sugar solutions versus a concentration gradient of the Cluster B EAA profile (1 to 1000 μM, n = 15 cages per group). There were no significant effects of EAAs on preferences (LME, day: F_1, 54_ = 0.0686, p = 0.79, EAA group: F_3, 54_ = 1.19, p = 0.32). Boxplots show acclimation (no choice) and test day (choice) preferences, with points representing individual cages. D: Total volume consumed from C (n = 15 cages per group). There were significant differences between acclimation (no choice) and test (choice) days (LME: day, F_1,54_ = 8.44, p = 0.0053; group, F_3,54_ = 0.103, p = 0.96; daygroup, F_3,54_ = 0.959, p = 0.42). Asterisks indicate significant post hoc comparisons (emmeans, Tukey adjustment).

The cluster B amino acid profile was tested over a concentration gradient from 1 to 1000 μM total EAAs. This cluster had the highest mean nectar concentrations of EAAs and included every EAA at concentrations between 0.44 µM (Trp) and 6.40 µM (Met). As observed in Figure 4B, the concentration of the cluster B EAA nectar profile did not affect the bees’ preference over sugar alone (Figure 4C). However, only at the highest concentration we tested were we able to observe that EAAs affected the total volume of all food consumed in the experiment (Figure 4D). There was no significant effect of any of the EAA mixtures on survival (SI Figure 2A-B).

### Bumblebees avoid drinking proline

Bees exhibited a weak preference for solutions containing 1 µM Pro, but in general, we found that bumblebees avoided drinking solutions containing Pro over concentrations from 10–100 µM (Figure 5A). None of the Pro solutions significantly affected survival over the duration of the assay (SI Figure 2C).

**Figure 5:**
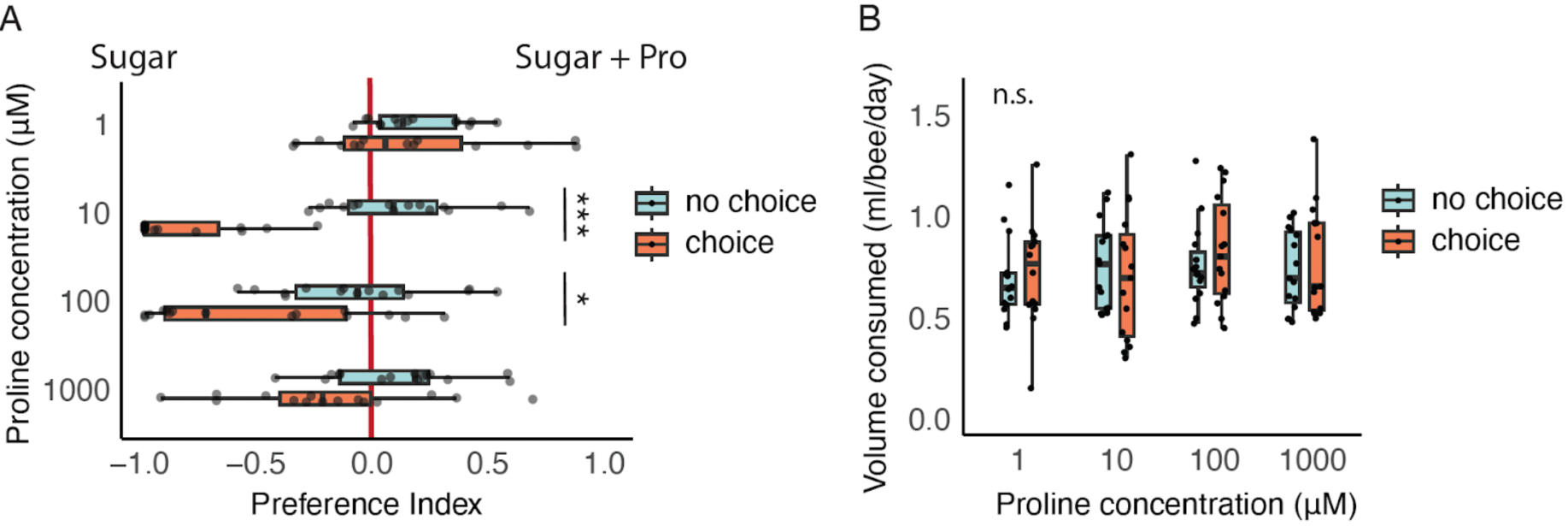
Bumblebee preferences for sugar solutions containing proline. A: Preference assay comparing the preferences of bumblebees to sugar mixtures with or without proline (1 – 1000 μM, n = 15 cages per group). Individual cages were tested with a no choice (sugar only) control day preceding the test day. Boxplots show median and interquartile range, with individual data points plotted for each cage. B: Total volume consumed (sugar solution and sugar + proline solution) on the no choice and choice days (n = 15 cages per group). Boxplots show median and interquartile range, with individual data points plotted for each cage.

## Discussion

The presence of AAs in nectar has been known for over 50 years (Baker & Baker, 1973), but whether they are a trait under selection by pollinators is poorly understood (Parachnowitsch *et al*., 2019). Here, we investigated the importance of AAs for pollinators by focussing on the relative proportions of the EAAs in nectar. We report evidence that while overall AA concentrations and the proportions of some AAs (Pro, Phe, and Met) of the nectar of UK plant species are phylogenetically conserved, variation in the complete profile of nectar EAAs is a much more divergent trait. In general, nectar EAAs increased feeding behaviour of bumblebees. Together this suggests that the blend of AAs in nectar may be a target for pollinator-driven selection.

Using the EAA profile (proportions of individual EAAs relative to the total AA concentration), the nectars clustered into six groups; in each case these profiles were characterised by one or two prominent EAAs. When we tested the nectar of these six groups on bumblebees, the bees did not prefer or avoid any of the solutions, indicating that they are unlikely to taste these AAs at micromolar concentrations. However, with some EAA profiles, the bees ate significantly more food overall. This suggests that low levels of EAAs in food provide a post-ingestive signal of value that drives feeding. This effect was found for four out of the six nectar EAA mixtures at nectar-relevant concentrations. In contrast, the NEAA, Pro, was generally avoided at concentrations > 1 μM but its presence did not affect the total amount of food eaten by the bees.

Unlike other floral traits such as colour or scent, there has been comparatively little research on the potential for pollinators to act as selective agents on nectar chemistry. For example, few other studies have quantified the degree of phylogenetic signal on nectar AAs (Gijbels *et al*., 2014). Interestingly, one exhaustive study of the 57 species in the family Balsaminaceae, found similar results to ours (Vandelook *et al*., 2019), reporting that the total AA concentration and the amount of Met both exhibited phylogenetic signal while the profile of AAs produced by a plant species did not. This supports our contention that it is the AA blend which may be of more importance to pollinators. However, they found that there was no evidence that AA nectar traits were associated with pollinator group (Vandelook *et al*., 2019). Contrastingly, a second study found no phylogenetic signal for total AA, EAA, or NEAA concentrations (Venjakob *et al*., 2022). However, while covering more plant families, this study involved fewer (34) species, which may limit detection of significant signal for any of the traits considered (Münkemüller *et al*., 2012).

The presence or absence of phylogenetic signal does not; however, directly imply low or high pollinator-driven selection (Abrahamczyk *et al*., 2017). A trait which shows high signal may be phylogenetically constrained, but equally, stabilising selection within closely related species could generate similar signal (Blomberg & Garland, 2002). Meanwhile low phylogenetic signal is indicative of selective pressure, but there are multiple agents (not just pollinators) which could be driving this (Parachnowitsch *et al*., 2019).

Nevertheless, although we found the concentration of nectar AAs may be phylogenetically constrained, our data show the profile of EAAs in nectar is a more labile trait, which could be driven by pollinator preferences. The behavioural experiments also suggest that EAA blends may be more important to pollinators than overall EAA concentrations. For example, both the EEA blends represented by cluster B and cluster D had Met as a significant proportion of the total EAA concentration. In contrast, while the blend of EAAs in cluster D stimulated feeding behaviour of bumblebees, the same was not true for cluster B. Furthermore, varying the total concentration of the EAA blend in cluster B across several orders of magnitude had little impact on feeding preference. Previous studies of AA preferences of pollinators have overwhelmingly focussed on individual AAs (Inouye & Waller, 1984; Carter *et al*., 2006; Bertazzini & Forlani, 2016; Hendriksma & Shafir, 2016; Bogo *et al*., 2024), while studies testing preferences for AA blends are few, particularly for bees (Stabler *et al*., 2015). Individual amino acids can either be aversive to bees or promote feeding (Inouye & Waller, 1984; Kim & Smith, 2000; Bertazzini & Forlani, 2016; Hendriksma & Shafir, 2016; Bogo *et al*., 2024), suggesting differential effects of individual AAs on reward valuation. Furthermore, not all AAs are detectable or discernible via chemotactile sensation in bumblebees (Ruedenauer *et al*., 2019) and wasps (Mattiacci *et al*., 2023). Taken together, this suggests that the relative blend of AAs is likely driving the feeding effects we observed with our EAA mixtures, rather than the presence or absence of individual AAs.

While including EAAs increased the amount of solution consumed, the bees did not avoid or prefer the solutions containing EAAs, regardless of their concentration, when they were tested over the μM range (Figure 4A). These data indicate that bees cannot taste low concentrations of EAAs, but that they have post-ingestive mechanisms for detecting them in food (Paoli *et al*., 2014; Stabler *et al*., 2015). EAAs such as the branched-chain AAs, Leu, Ile, and Val, are known to drive feeding post-ingestively in mammals (Solon-Biet *et al*., 2019), insects (Piper *et al*., 2017), and in bees (Stabler *et al*., 2015). Relatively high concentrations of free EAAs can be aversive or even toxic to bees (Paoli *et al*., 2014). For this reason, we predict that although the concentrations of these EAAs are low in nectar, they are in a range that is not aversive but still valuable to pollinators as a source of essential nutrients. We expect this post-ingestive effect takes a longer time (s) for the brain to process than taste cues (ms) (Simcock *et al*., 2018).

Two individual EAAs of note were Met and Phe. Across four of the six nectar clusters one of these two compounds formed a dominant component of the EAA blend. This occurs repeatedly across all the families we sampled. This pattern was particularly obvious for Phe, as eight out of the 13 families sampled had at least one species with Phe as the dominant EAA. Phe has previously been reported as a phagostimulant for honeybees (Hendriksma & Shafir, 2016) and other insects (Dethier, 1976). It is also a dominant feature of nectar of many bee-visited plants, especially in the Lamiaceae (Petanidou *et al*., 2006).

Proportionally high concentrations of Met in nectar occurred in seven out of the eight Plantaginaceae species in our dataset. Met is particularly important as a dietary component for egg production in insects (Grandison *et al*., 2009; Lee *et al*., 2014). However, high concentrations of Met can be toxic and reduce lifespan (Manoukas, 1981; Grandison *et al*., 2009; Lee *et al*., 2014). Dietary Met could be particularly important for bees: free Met is needed for genome methylation especially during development, and methylation is a mechanism important for caste differentiation and other processes in eusocial bees, as ∼35% of the bee genome is methylated (Foret *et al*., 2009). This is in contrast to other insects like Drosophila where only ∼ 1% is methylated (Takayama *et al*., 2014). Owing to the low absolute concentrations of any AAs in nectar, any implications for bee nutrition are likely to be less important than effects on feeding behaviour. This is because bees rely on pollen for most of their protein and AA requirements (Wright *et al*., 2018).

Previous work has found that, often, one or two AAs occur at much higher concentrations than the remaining AA complement of nectar [e.g., Phe, (Petanidou *et al*., 2006), Ala (Göttlinger & Lohaus, 2024), and Pro (Kaczorowski *et al*., 2005)]. By comparison with all the other AAs in our dataset, the NEAA Pro was the most common. Pro was present in 87% of species at > 0.001 μM and highly abundant, for example up to 11 mM in the bumblebee pollinated yellow iris, *Iris pseudocorus*. Pro is clearly important for pollinators: it forms, by far, the dominant component of haemolymph AAs in honeybees (Crailsheim & Leonhard, 1997) and bumblebees (Stabler *et al*., 2015). It also can power insect flight (Bursell 1975); though in bees the importance of this role is disputed (Micheu *et al*., 2000; Carter *et al*., 2006; Syromyatnikov *et al*., 2013; Stec *et al*., 2021). Interestingly, in our behavioural experiments, the bees were very sensitive to the presence of Pro in the nectar solutions, as they detected and avoided quite low (10 μM) concentrations of Pro. Curiously, their aversion to Pro was reduced as its concentration increased from 10 μM to 1000 μM in nectar.

Plants are likely to have mechanisms for the active regulation of the quantities of AAs in nectar, but only a few studies have tested this. For example, in bromeliads, the AA concentrations in leaves and nectaries were much greater than in nectar (Göttlinger & Lohaus, 2022). This regulation may not be complete though, as several studies have shown that experimental addition of fertiliser leads to higher nectar AA concentrations (Gardener & Gillman, 2001b; Gijbels *et al*., 2015a). However, regulation of AA composition at the level of the nectaries also appears to extend to individual AAs or the AA profile (Lohaus & Schwerdtfeger, 2014; Bertazzini & Forlani, 2016; Göttlinger & Lohaus, 2024). Particularly striking is another example of two groups of species from the bromeliad genus *Pitcarnia*. Both groups have high Ala concentrations in phloem tissue, but species in one group have high Ala concentrations in the nectar as well, whereas the others do not (Göttlinger & Lohaus, 2024).

Our data show indicate that pollinators do play a role in shaping the presence and concentration of AAs in nectar. While it is obvious that carbohydrate composition correlates with pollinator guild (Abrahamczyk et al., 2017 and others), such evidence is currently lacking for AAs. Our data suggest that EAA profile is a nectar trait potentially under selection by pollinators and that the EAA profile is more important than EAA concentration in its effect on pollinator feeding behaviour. We suggest future research further investigates the role of EAA profiles in determining the extent to which nectar AAs drive plant-pollinator coevolution.

## Supporting information

Supplemental_Information_main

## Acknowledgements

This research was funded by BBSRC grants (BB/S000402/1, BBI000968/1) and a Leverhulme Trust grant (RPG-2020-393) awarded to GAW, as well as a Royal Society Newton International Fellowship (NIF\R1\191468) awarded to RHP.

## Author contributions

R.H.P. and G.A.W. conceptualised the study. R.H.P. and G.A.W. developed the methodology. Formal analysis was conducted by R.H.P. and A.E.MR. The investigation was carried out by E.F.P., R.H.P., K.W., and A.E.MR. Data curation was performed by R.H.P. and A.E.MR. Visualisations were prepared by R.H.P. R.H.P., J.G.P., and G.A.W. wrote the original draft of the manuscript. All authors contributed to reviewing and editing the manuscript. Funding was acquired by G.A.W.

## Data availability

All data and code are available for review at: https://drive.google.com/drive/folders/1V2AtNXChfT804Q6xL4a-i5cFEytaX8qc?usp=sharing

Public GitHub repository will be made available upon publication.

