## Supplemental_Information_main for "Do pollinators play a role in shaping the essential amino acids found in nectar?"

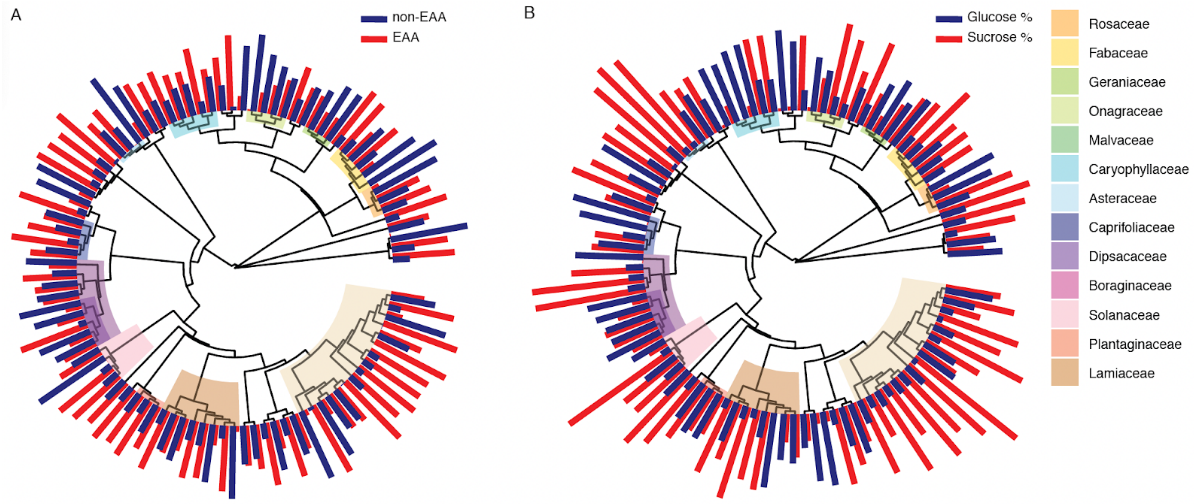


**SI Figure 1:** Phylogenetic trees constructed with 96 species from our nectar dataset. Red and blue bars display the total concentrations of NEAAs and EAAs (A) and the proportions of glucose and sucrose (B) across the dataset, with families highlighted on each tree where the number of species is >2.

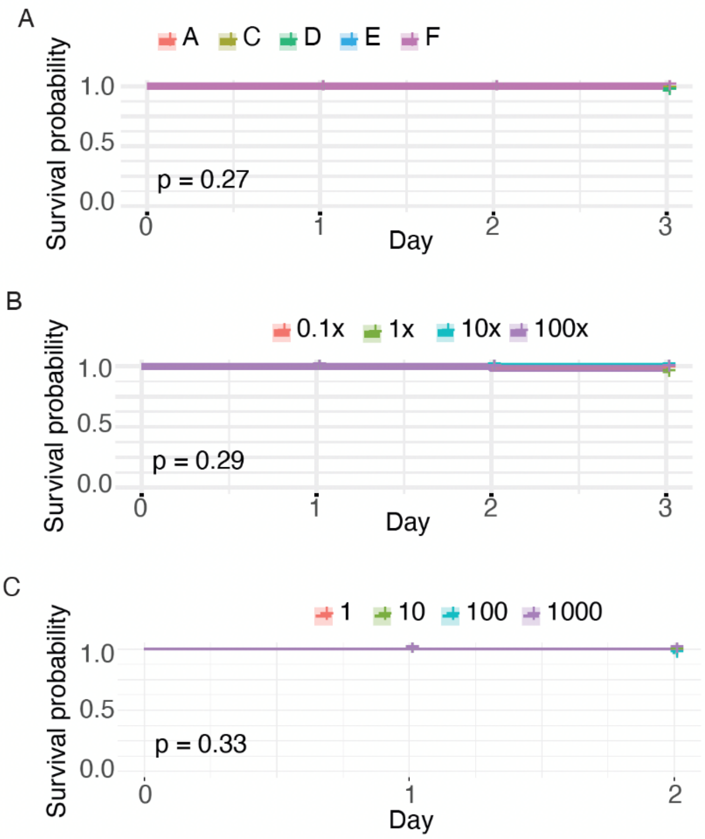


**SI Figure 2:** Kaplan-Meier mortality curves for the three bumblebee preference assays: nectar relevant concentrations of cluster A, C, D, E, and F nectar (A), a concentration gradient of cluster B nectar profile (B) and the proline preference assay (C). There was minimal mortality across any of the assays, and no significant difference between groups (n = 75 bees per group). Exact p-values shown on each plot.

**SI Table 1: Summary of the mean and standard deviation of concentrations of sugars and amino acids across samples for each species.**

*** Table is attached (too large to display) ***

**SI Table 2: Results from phylogenetic analysis using Blomberg's K and Pagel's Lambda.** The EAA profiles (t-SNE dimensions), proportions of individual EAAs and proline, total concentrations of EAAs, NEAAs, and sugars, and proportions of sugars were tested. Significant p-values are highlighted in orange.

| Nectar variable | Blomberg's K | | Pagel's Lambda | |
|---|---|---|---|---|
| | stat | p-value | stat | p-value |
| t-SNE Dim 1 | 0.066 | 0.209 | 0.000 | 1.000 |
| t-SNE Dim 2 | 0.051 | 0.682 | 0.000 | 1.000 |
| His % | 0.051 | 0.566 | 0.000 | 1.000 |
| Thr % | 0.031 | 0.978 | 0.000 | 1.000 |
| Arg % | 0.056 | 0.475 | 0.000 | 1.000 |
| Val % | 0.048 | 0.771 | 0.000 | 1.000 |
| Met % | 0.099 | 0.060 | 0.577 | 0.042 |
| Trp % | 0.034 | 0.836 | 0.000 | 1.000 |
| Phe % | 0.094 | 0.038 | 0.044 | 0.530 |
| Ile % | 0.062 | 0.336 | 0.000 | 1.000 |
| Leu % | 0.042 | 0.744 | 0.000 | 1.000 |
| Lys % | 0.046 | 0.798 | 0.000 | 1.000 |
| Pro % | 0.100 | 0.018 | 0.035 | 0.669 |
| EAA (μM) | 0.440 | 0.009 | 0.913 | 0.001 |
| NEAA (μM) | 0.137 | 0.018 | 0.346 | 0.384 |
| Sugars (M) | 0.073 | 0.129 | 0.000 | 1.000 |
| Sucrose % | 0.084 | 0.024 | 0.026 | 0.695 |
| Fructose % | 0.071 | 0.136 | 0.000 | 1.000 |
| Glucose % | 0.080 | 0.036 | 0.067 | 0.420 |
